# Amyloid β oligomers dysregulate oligodendrocyte differentiation and myelination via PKC in the zebrafish spinal cord

**DOI:** 10.1101/2024.06.10.598246

**Authors:** U. Balantzategi, A. Gaminde-Blasco, L. Bayón-Cordero, M. V. Sánchez-Gómez, J. L. Zugaza, B. Appel, E Alberdi

## Abstract

Amyloid β oligomers (Aβo) have been proposed as candidates to induce oligodendrocyte (OL) and myelin dysfunctions in early stages of Alzheimer’s disease (AD) pathology. Nevertheless, little is known about how Aβo affect OL differentiation and myelination *in vivo*, and the underlying molecular mechanisms. In this study, we explored the effects of a brain intraventricular injection of Aβo on OLs and myelin in the developing spinal cord of zebrafish larvae. Using quantitative fluorescent *in situ* RNA hybridization assays, we demonstrated that Aβo altered *myrf* and *mbp* mRNA levels and the regional distribution of *mbp* during larval development, suggesting an early differentiation of OLs. Through live imaging of Tg(*myrf*:mScarlet) and Tg(*mbp*:tagRFP) zebrafish lines, both crossed with Tg(*olig2*:EGFP), we found that Aβo increased the number of *myrf*^+^ and *mbp*^+^ OLs in the dorsal spinal cord at 72 hpf and 5 dpf, respectively, without affecting total cell numbers. Furthermore, Aβo also increased the number of myelin sheaths per OL and the number of myelinated axons in the dorsal spinal cord compared to vehicle-injected control animals. Interestingly, the treatment of Aβo-injected zebrafish with the pan-PKC inhibitor Gö6983 restored the aforementioned alterations in OLs and myelin to control levels. Altogether, not only do we demonstrate that Aβo induce a precocious oligodendroglial differentiation leading to dysregulated myelination, but we also identified PKC as a key player in Aβo-induced pathology.

## Introduction

Oligodendrocytes (OLs) are specialized glial cells responsible for forming myelin sheaths around axons in the central nervous system (CNS). This myelination is essential for the rapid propagation of electrical signals and overall neuronal connectivity (Baumann & Pham-Dinh, 2001). Dysfunction of OLs and myelin are among the early pathological events observed in Alzheimer’s disease (AD), a chronic neurodegenerative condition primarily characterized by cognitive decline and memory loss (Desai et al., 2009; Roher et al., 2002). Although there is strong evidence connecting OL impairment with the onset of neurodegeneration in AD, the underlying causes of OL dysfunction in this disorder are poorly understood. One of the major pathological hallmarks of AD is the accumulation of amyloid-β oligomers (Aβo), which are thought to play a critical role in the disruption of OL differentiation and myelination processes (Mitew et al., 2010). Only a few studies have investigated the role of OLs in AD and proposed Aβ as a candidate for promoting white matter dysfunction (Dean et al., 2017; Selkoe & Hardy, 2016), but the mechanisms remain unclear.

Protein kinase C (PKC) is a family of kinases that play essential roles in various physiological processes to maintain cell homeostasis, including cell cycle regulation (Nelson & Alkon, 2009), differentiation (Cavaliere et al., 2013; Damato et al., 2021), and apoptosis. Interestingly, several studies have shown that PKC activation can influence OL differentiation, inducing the expression of myelin-related genes and promoting myelination (Asotra & MacKlin, 1993; Swire et al., 2019). Moreover, PKC is involved in the pathogenesis of a wide range of disorders, including cancer (Garg et al., 2014) and neurological disorders such as AD (Alfonso et al., 2016; Newton, 2010). Furthermore, previous investigations have reported increased PKC activity in response to Aβ in both astrocytes (Abramov & Duchen, 2005; Wyssenbach et al., 2016) and neurons (Manterola et al., 2013; Ortiz-Sanz et al., 2022). However, Aβ-induced PKC activation has not been explored for OLs.

Zebrafish (*Danio rerio*) are increasingly recognized as a valuable model organism for studying OLs and myelin *in vivo*, since its embryos and larvae are transparent, allowing for real-time imaging of myelination and OL dynamics (Kimmel et al., 1995). In addition, zebrafish have a high degree of genetic and physiological homology with humans, particularly in the CNS, making them a relevant model for human neurological diseases (Howe et al., 2013; Lieschke & Currie, 2007). Therefore, they have become a valuable tool for conducting pharmacological studies (Cassar et al., 2020).

In this study, we investigated whether intracerebroventricular (ICV) injection of Aβo into zebrafish larvae affects OL differentiation and myelination *in vivo*, and whether PKC is involved in the underlying signaling pathway. Our results indicated that Aβo alter *myrf* and *mbp* mRNA levels and the regional distribution of *mbp* in the developing spinal cord of zebrafish larvae. Live imaging revealed that Aβo increased the number of *myrf*^+^ and *mbp*^+^ OLs in the dorsal spinal cord without affecting the total cell number, suggesting premature differentiation and migration of OLs. Additionally, Aβo increased the number of myelin sheaths per OL and the number of myelinated axons in the dorsal spinal cord without affecting the sheath length. Treatment with the pan-PKC inhibitor Gö6983 reversed the effects of Aβo to physiological levels, confirming that Aβo dysregulated OLs and myelin through PKC function. These results provide new insights into the *in vivo* effects of Aβo on OLs and highlight PKC as a promising therapeutic target in AD.

## Material and methods

### Zebrafish

Nontransgenic embryos were obtained through crosses of male and female zebrafish from the AB strain. Embryos were raised at 28.5 °C in E3 media (5 mM NaCl, 0.17 mM KCl, 0.33 mM CaCl, 0.33 mM MgSO4 (pH 7.4), supplemented with sodium bicarbonate). Larvae were staged according to hours or days post-fertilization (hpf/dpf), and selected based on criteria ensuring good health and normal developmental patterns. For cell count experiments, the previously established zebrafish lines *Tg(olig2:EGFP)^vu12^*, *Tg(myrf:mScarlet)^co66^*, and *Tg(mbpa:tagRFPT)^co25^*were used.

### Preparation of amyloid β oligomers (Aβo)

Oligomeric amyloid-β (Aβ1-42) was prepared as previously described (Dahlgren et al., 2002). Briefly, Aβ1-42 (Bachem, Germany) was initially dissolved to a concentration of 1 mM in hexafluoroisopropanol (Sigma-Aldrich), which was next totally removed under vacuum using a speed vac system. The resulting peptide film was stored desiccated at -80 °C. For the aggregation protocol, the peptide was resuspended in dry dimethylsulfoxide (DMSO; Sigma-Aldrich) to achieve a concentration of 5 mM, and Hams F-12 (PromoCell) was then added to adjust the peptide to a final concentration of 100 μM (the vehicle consisted of DMSO in Hams F-12). Oligomer formation was induced by incubating the peptide solution for 24 h at 4 °C and confirmed by polyacrylamide gel electrophoresis (SDS-PAGE) followed by Coomassie Brilliant Blue R-250 (Bio-Rad) staining.

Fluorescently labeled Aβ1-42 was prepared as previously described (Jungbauer et al., 2009). After 24 h incubation at 4 °C, labeling was performed using the Alexa Fluor^TM^ 488 Microscale Protein Labeling Kit (#A30006, Invitrogen) according to manufactureŕs instructions. Briefly, 50 µl of 100 µM Aβo solution was adjusted to pH 9 with 5 µl of 1 M NaHCO3, followed by addition of 4 µl of the H2O solubilized reactive dye. Incubation at room temperature was performed for 15 min followed by immediate addition of 55 µl of the labeling reaction mixture onto a spin column packed with a 425 µl slurry of Biogel P-6 resin for removal of unincorporated dye. The resulting eluent (pH 7.4) was stored for up to 2 days at 4 °C or used immediately for injections.

### Intracerebroventricular amyloid β injection in zebrafish larvae

At 24 hours post fertilization (hpf), all zebrafish embryos had their chorions removed for the brain ventricle injection procedure following previously established protocols (Gutzman & Sive, 2009; Nery et al., 2014). Embryos were anesthetized with Tricaine (Sigma-Aldrich) and immobilized in wells on 2%-agar coated dishes under the stereomicroscope so that the brain ventricle was exposed. An injection needle was carefully positioned on the roof plate of the hindbrain, and 5–10 nl of a fresh injection buffer containing 10% either Aβo (10 µM) or vehicle, and 10% Phenol Red in 0.4 M KCl in nuclease-free water was microinjected. Subsequently, each treatment group was separated into clean plates, and the zebrafish were returned to the incubator and allowed to grow until the days of the experiments.

### RNA *in situ* hybridization

To identify potential Aβo-induced alterations in OL differentiation- and myelination-related gene expression *in vivo*, we conducted fluorescent RNA *in situ* hybridization (FISH) assays in the spinal cords of zebrafish larvae injected with either vehicle or Aβo (10 µM). FISH was performed using the RNAScope Multiplex Fluorescent V2 Assay Kit (Advanced Cell Diagnostics). Zebrafish larvae at 48 hpf, 72 hpf, and 5 dpf were fixed in 4% PFA in PBS, gently rocking O/N at 4 °C. Subsequently, samples were embedded in 1.5% agar with 30% sucrose, followed by immersion in 30% sucrose overnight. The blocks were frozen on dry ice, and 15 μm-thick transverse sections were obtained using a cryostat microtome and collected on polarized slides. FISH was performed according to the manufacturer’s instructions, with the following modification: slides were covered with Parafilm for all 40 °C incubations to maintain moisture and disperse reagents across the sections. The zebrafish *mbpa*, *myrf*, and *sox10* transcript probes were designed and synthesized by the manufacturer, and used at 1:50 dilutions, except for *mbpa*, which was used undiluted. The transcripts were fluorescently labeled by TSA-based Opal fluorophores Opal520 (1:1500), Opal570 (1:500) and Opal650 (1:1500) using the Opal 7 Kit (PerkinElmer).

Images were acquired on a Zeiss Cell Observer 2D 25 Spinning Disk confocal system (Carl Zeiss Microscopy) with a 40X oil-immersion objective. 15 z-stack tiles (z step = 0.5 µm) of 5 sections of the spinal cord were acquired from each animal. The total area occupied by each probe was determined using ImageJ/Fiji.

### *In vivo* myelin sheath visualization

To visualize myelin sheaths *in vivo*, the mbpa:EGFP-CAAX construct was transiently expressed by microinjection into 1-cell stage nontransgenic zebrafish embryos. Next, zebrafish larvae were ICV injected with either vehicle or Aβo (10 µM) at 24 hpf, and treated by bath immersion with or without Gö6983 (500 nM, Tocris) at 72 hpf. At 5 dpf, zebrafish larvae were anesthetized with Tricaine (Sigma-Aldrich) and embedded laterally in 1% low-melt agarose containing 0.4% Tricaine for immobilization on a glass bottom dish.

Images were acquired using a Zeiss Cell Observer 2D 25 Spinning Disk confocal system (Carl Zeiss Microscopy) with a 40X water-immersion objective. Images were collected from 1-3 field of view from each animal in 40 z-stack tiles (z-step = 0.28 μm). Subsequent analysis of sheath length and total number of sheaths per OLs was performed using Imaris software v9.9.1 (Oxford Instruments), in a blind mode.

### Electron microscopy (EM)

At 8 dpf, vehicle- or Aβo-injected (10 µM) zebrafish larvae were anesthesized with Tricaine and fixed in a solution consisting of 2% glutaraldehyde and 4% PFA in 0.1 M sodium cacodylate (pH 7.4), for at least 5 days at 4°C. The tissue was then sagittally sectioned using a Leica VT 1200S vibrating blade microtome (Leica) to obtain 200 μm-thick sections. These tissue sections were post-fixed in 2% OsO4, dehydrated in ethanol and propylene oxide, and embedded in EPON (Serva) for 24 h at 60 °C. Ultrathin sections (50 nm thickness) were obtained using a Leica Ultracut S ultramicrotome (Leica) and contrasted for 30 min with 4% uranyl acetate and 6 min with lead citrate.

Electron micrographs were captured using a Zeiss EM900 electron microscope (Carl Zeiss Microscopy), and the number of dorsal myelinated axons were quantified using ImageJ/Fiji.

### Statistical analysis

All data are presented as mean ± S.E.M (standard error of the mean), with sample size indicated in the figures by dots. Statistical analyses were performed using absolute values. GraphPad Prism 8.2.1 software was utilized, applying unpaired two-tailed Student’s t-test for comparing two experimental groups, and one-way or two-way analysis of variance (ANOVA) followed by Tukey’s and Sidak’s *post-hoc* tests for multiple comparisons. Results from independent animals were considered as biological replicates (n ≥ 5). Statistical significance was represented as p<0.05 (*), p<0.01 (**), and p<0.001 (***).

## Results

Emerging evidence suggests that alterations in oligodendrogenesis and disruptions in myelin integrity are key early events in AD pathology (Ferreira et al., 2020; Wu et al., 2017). Here, we hypothesized that Aβo may significantly influence these processes. Consequently, our main goal was to investigate the potential effects of ICV injection of Aβo on OL differentiation and myelination *in vivo* in zebrafish larvae.

First, proper Aβ oligomerization was confirmed using in-gel Coomassie blue staining (**Figure 1A**). In addition, the efficiency of injections was assessed by injecting fluorescently labeled dextran (**Supplementary figure 1**) and Aβo (**Figure 1B**) into 24 hpf zebrafish larvae. We observed a widespread distribution of the injection mixture throughout the brain and spinal cord of the larvae. Therefore, ICV injections were considered adequate to proceed with the proposed experimental procedures.

**Figure 1.**
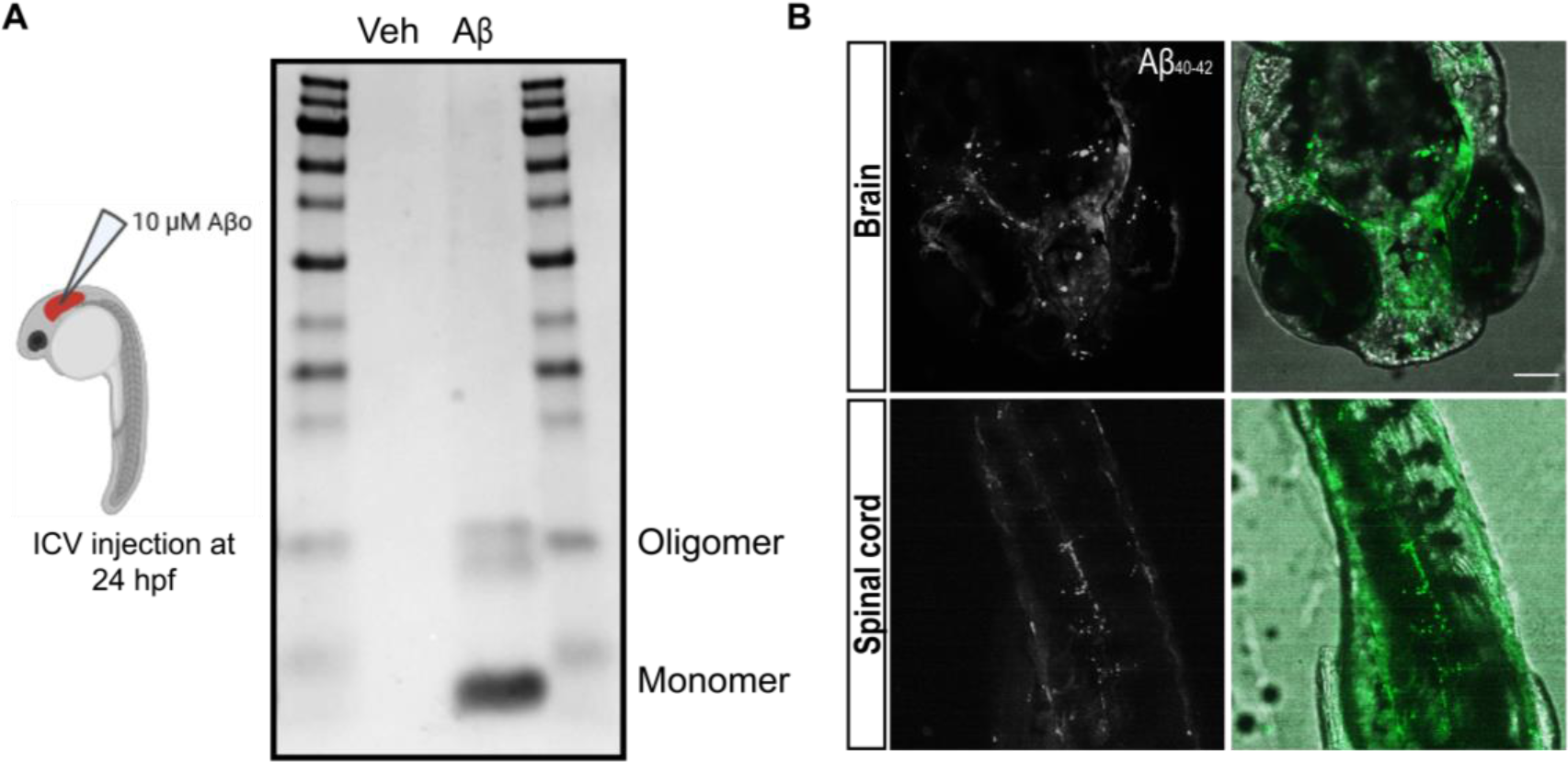
Intracerebroventricular Aβo injection into zebrafish larvae. (**A**) Schematic representation of the experimental approach, and the detection of Aβ species (monomers and oligomers) in the injection mixtures of vehicle and Aβ using Coomassie blue staining. (**B**) Representative fluorescent images of labeled-Aβo diffusion into the brain and spinal cord of 24 hpf zebrafish larvae following ICV injection. Scale bar = 50 µm.

### Aβo induce early oligodendrocyte differentiation and myelination in zebrafish

To identify Aβo-induced alterations in OL differentiation and myelination, we conducted fluorescent RNA *in situ* hybridization (FISH) assays following Aβo or vehicle ICV injections. The expression of three key genes associated with different developmental stages of the OL lineage was assessed: *sox10*, a canonical marker of OL lineage*; myrf,* expressed in differentiating OLs (Hornig et al., 2013); and *mbpa* (*mbp*), a zebrafish orthologue of murine and human *MBP*, expressed in myelinating OLs (**Figure 2A**).

**Figure 2.**
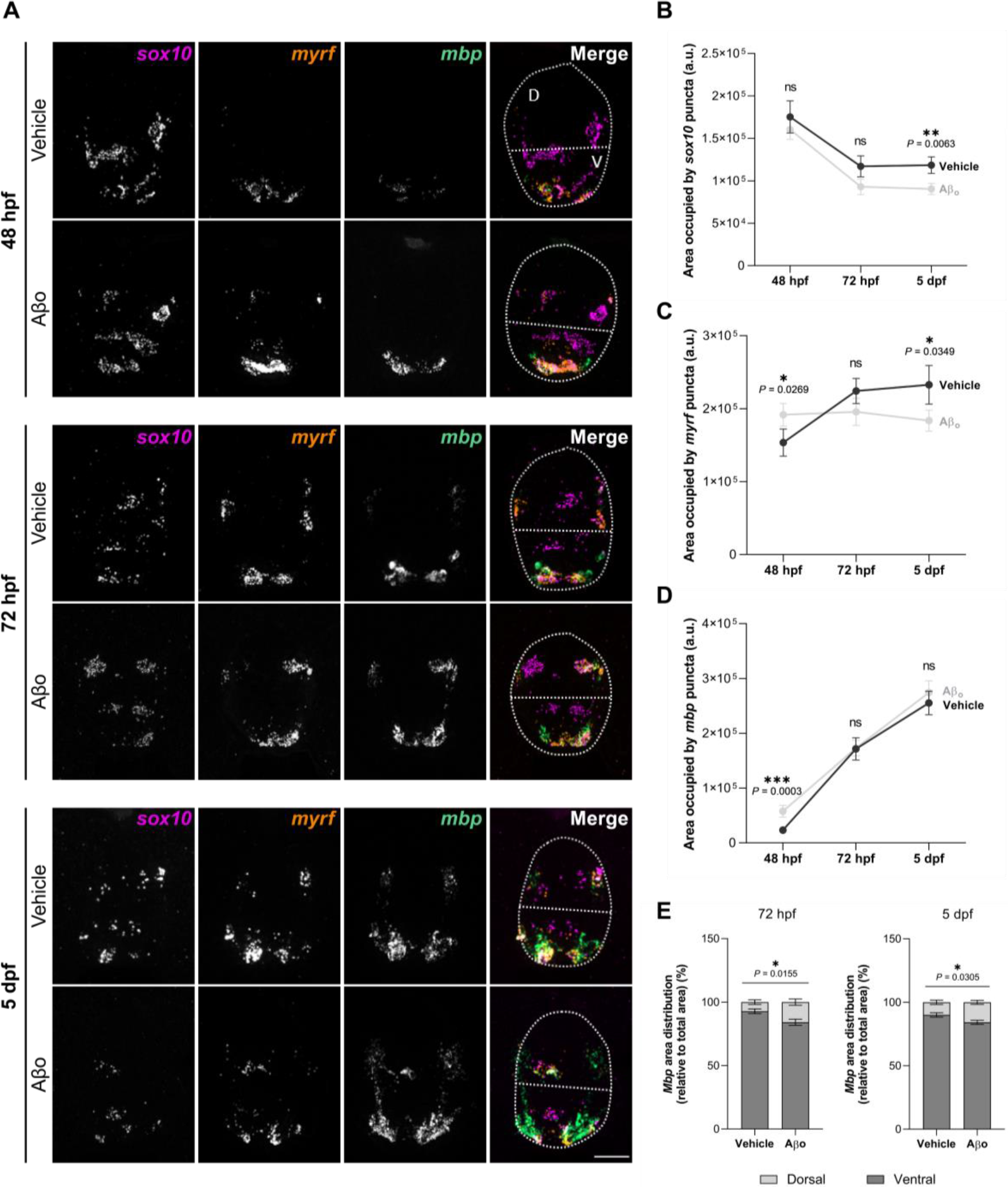
Aβo-injected zebrafish larvae exhibit alterations in oligodendroglial lineage mRNA levels and regional distribution in the developing spinal cord. Zebrafish larvae were injected with Aβo or its vehicle at 24 hpf, and FISH assays were performed for *sox10*, *myrf* and *mbp* at 48 hpf, 72 hpf and 5 dpf. (**A**) Representative confocal images of spinal cord cross-sections depicting mRNA expression fluorescence. (**B**, **C**, **D**) RNAscope analysis representing the area occupied by each mRNA in the developing spinal cord. (**E**) Quantification of *mbp* mRNA distribution in ventral and dorsal regions. Scale bar = 20 µm. Data are represented as means ± S.E.M, p*<0.05, p**<0.01, p***<0.001 compared to vehicle-injected zebrafish larvae. Statistical significance was drawn by two-tailed nested t-test and two-way ANOVA followed by Sidak’s *post-hoc* test. n = 9 - 11 larvae per condition and time-point.

The expression of *sox10* decreased during the initial days of larvae development, particularly in Aβo-treated zebrafish (1.18 ± 0.1x10^5^ a.u. in vehicle-treated larvae at 5 dpf vs 0.9 ± 0.07x10^5^ a.u. in Aβo-treated larvae) (**Figure 2B**). On the other hand, interestingly, *myrf* expression was significantly increased by ∼25% at 48 hpf in the presence of Aβo (1.54 ± 0.19x10^5^ a.u. vehicle-treated vs 1.92 ± 0.16x10^5^ a.u. Aβo-treated larvae). However, while *myrf* mRNA showed an expected increasing expression throughout development in control larvae, Aβo induced a more stable expression of *myrf* during time, resulting in a ∼21% reduced expression at 5 dpf compared to vehicle-injected larvae (2.33 ± 0.27x10^5^ a.u. vehicle vs 1.84 ± 0.15x10^5^ a.u. Aβo) (**Figure 2C**). Consistent with increased *myrf* levels at 48 hpf, there was a notable increase of ∼147% in *mbp* expression levels at that same time-point in presence of Aβo (2.33 ± 0.33x10^4^ a.u. vehicle vs 5.77 ± 1.08x10^4^ a.u. Aβo). However, this early boost of the myelin-related mRNA was normalized by 72 hpf, and remained at control levels throughout development (**Figure 2D**).

In zebrafish spinal cord development, the majority of OPCs originate from progenitor cells in the pMN domain located in the ventral spinal cord, and they then migrate towards the dorsal zone, where they undergo proliferation and differentiation into OLs (Lee et al., 2010). Consequently, not only differentiation-related gene expression but also OPC migration is crucial for proper OL differentiation and CNS development. Here, we observed a significant shift in *mbp* expression towards the dorsal area in Aβo-injected larvae both at 72 hpf (only 7.2 ± 1.82% of the total area occupied by *mbp* was dorsal in vehicle-injected larvae, compared to 15.9 ± 2.45% in Aβo-injected larvae) and 5 dpf (vehicle 9.95 ± 1.68% vs Aβo 15.77 ± 1.56%) (**Figure 2E, Supplementary figure 2A**). However, we did not observed changes in the regional distribution of *myrf* and *sox10* (**Supplementary figure 2B, C**).

Overall, these results suggested that Aβo might induce a precocious OL differentiation and myelination, by altering differentiation- and myelination-related mRNA expression and regional distribution during larval development.

### Aβo-induced deregulation of oligodendrocyte differentiation and maturation timing is PKC-dependent

Considering previous studies indicating increased PKC activity in response to Aβ in both astrocytes (Abramov & Duchen, 2005; Wyssenbach et al., 2016) and neurons (Manterola et al., 2013; Ortiz-Sanz et al., 2022), we next investigated the implication of PKC in Aβo-induced effects in OLs. Increased phosphorylation levels of PKC were found in primary cultured OLs treated with Aβo for 3 or 24 h assessed via western blot analysis (141.3 ± 10.09% after 3 h of Aβo treatment and 144.5 ± 12.38% after 24 h of Aβo treatment, compared to 100% in control cells)y, indicating Aβo-induced PKC activation in OLs (**Supplementary figure 3A, B**).

Thereafter, to explore Aβo-triggered alterations in OL differentiation more in depth, we conducted *in vivo* studies using stable transgenic zebrafish lines. Specifically, we crossed *Tg(olig2:EGFP)^vu12^*, a reporter of pMN lineage cells, with *Tg(myrf:mScarlet)^co66^*, a reporter of differentiating OLs. In addition to ICV injection of Aβo or vehicle at 24 hpf, some larvae were also treated with the pan-PKC inhibitor Gö6983 at 48 hpf (**Figure 3A**). Combination of Aβo and the drug did not show any noticeable significant toxicity for the larvae (**Figure 3B**).

**Figure 3.**
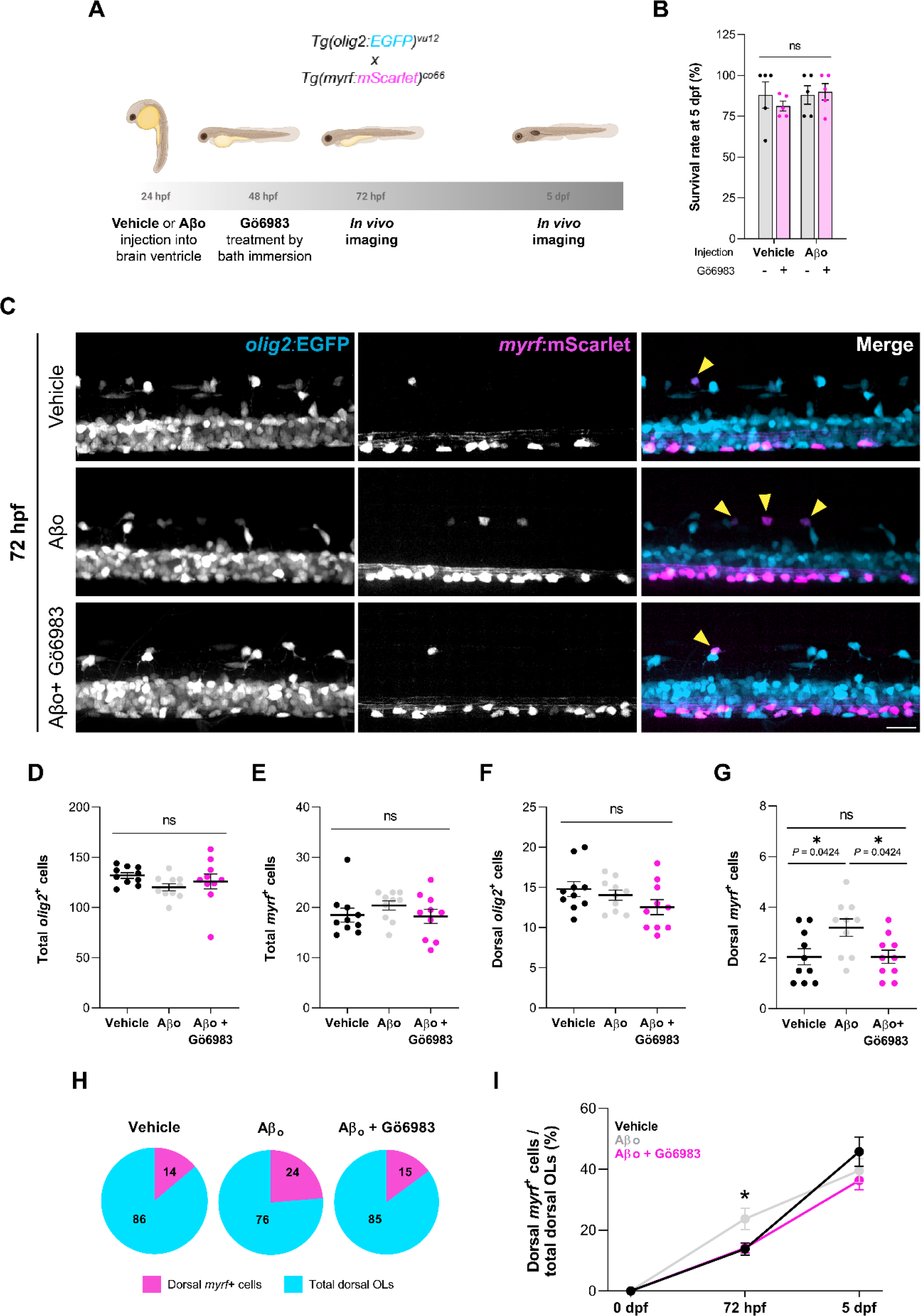
Aβo induce early oligodendrocyte differentiation through PKC, without affecting total cell numbers. (**A**) Transgenic zebrafish stably expressing *olig2*:EGFP and *myrf*:mScarlet were injected with Aβo or its vehicle at 24 hpf, and treated with Gö6983 (500 nM) at 48 hpf by bath immersion. Live imaging was performed at 72 hpf and 5 dpf. (**B**) Survival rate was measured at 5 dpf. (**C**) Representative lateral fluorescence images of the spinal cord in live transgenic larvae at 72 hpf. Yellow arrowheads indicate *olig2*^+^*myrf*^+^ cells. (**D**) Graphs comparing the number of total *olig2*^+^ cells, (**E**) total *myrf*^+^ cells, (**F**) dorsal *olig2*^+^ cells, and (**G**) dorsal *myrf*^+^ cells. (**H**) Pie charts showing the ratio of differentiating dorsal OLs (percentage of dorsal *myrf*^+^ cells among dorsal *olig2*^+^ cells) at 72 hpf for each condition. (**I**) Graph showing the progression of differentiating dorsal OLs during development. Scale bar = 20 μm. Data indicate means ± S.E.M, and dots represent individual larvae. *p<0.05; Statistical significance was determined by ordinary one-way ANOVA followed by Sidak’s *post-hoc* and mixed-effects analysis followed by Tukey’s *post-hoc.* n^72^ ^hpf^ = 10 larvae per condition; n^5^ ^dpf^ = 7-9 larvae per condition

At 72 hpf, no significant differences were observed in the total number of *olig2*^+^ (**Figure 3C, D**) or *myrf*^+^ cells (**Figure 3E**) across treatments. Nevertheless, when focusing on the dorsal spinal cord, while *olig2*^+^ cell counts remained consistent across all treatment groups (**Figure 3F**), we detected a notable increase in dorsal *myrf*^+^ cells in larvae injected with Aβo compared to those receiving vehicle injections. Interestingly, treatment with Gö6983 effectively attenuated Aβo-induced excess of *myrf*^+^ cells numbers to control values (vehicle 2.05 ± 0.44 cells vs Aβo 3.2 ± 0.39 cells vs Aβo + Gö6983 2.05 ± 0.49 cells) (**Figure 3G**). Indeed, Aβo-injected larvae exhibited an augmented proportion of differentiating OLs in the dorsal spinal cord at this larval stage: while only 14% of dorsal OLs were *myrf*^+^ in vehicle-treated larvae, this ratio increased to 24% following Aβo treatment and was maintained at 15% when Aβo-injected larvae were treated with Gö6983 (**Figure 3H**).

No significant changes were observed in differentiating *myrf*^+^ OL numbers at 5 dpf (**Supplementary figure 4**). Therefore, the sequential study of OL differentiation in this transgenic line suggested that Aβo induce an initial increased premature dorsal OL differentiation around 72 hpf that returns to normal levels by 5 dpf. This effect was reversed by PKC inhibition (**Figure 3I**).

The above data suggest that a higher proportion of OPCs undergo differentiation into mature myelinating OLs in Aβo-injected larvae. To directly test this possibility, we employed a similar approach as previously described, now crossing *Tg(olig2:EGFP)^vu12^*with *Tg(mbpa:tagRFPT)^co25^*, a reporter line for mature myelinating OLs (**Figure 4A**). Since changes in *myrf* were observed at 72 hpf and OL maturation occurs later in time, we focused our analysis on 5 dpf to examine mature myelinating OLs.

**Figure 4.**
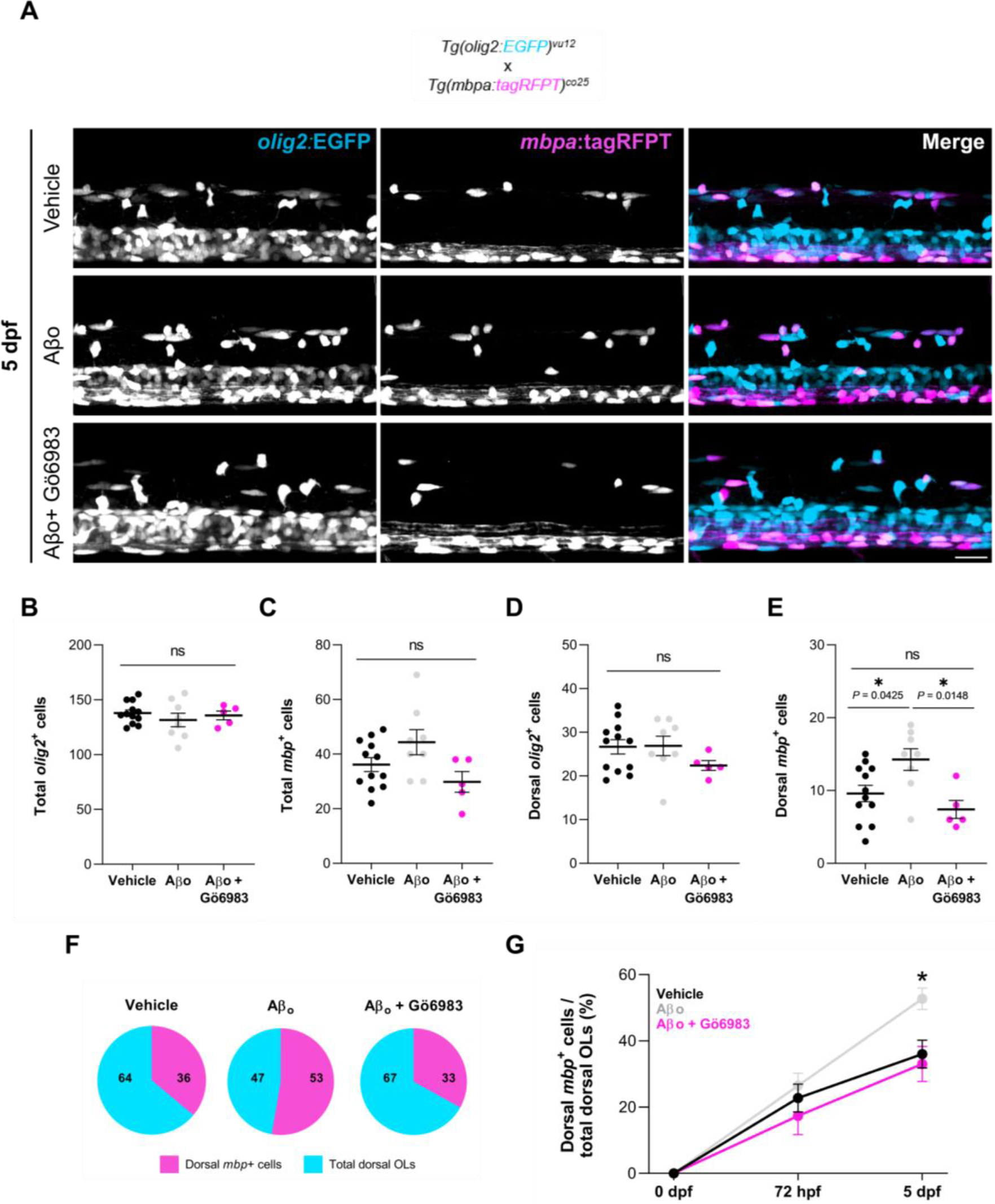
Aβo promote oligodendrocyte maturation via PKC, without changing total cell numbers. (**A**) Representative lateral images of the spinal cord of live transgenic larvae stably expressing *olig2*:EGFP and *mbpa*:tagRFPT, at 5 dpf. (**B**) Graphs comparing the quantity of total *olig2*^+^ cells, (**C**) total *mbp*^+^ cells, (**D**) dorsal *olig2*^+^ cells, and (**E**) dorsal *mbp*^+^ cells. (**F**) Pie charts showing the ratio of mature myelinating dorsal OLs (percentage of dorsal *mbp*^+^ cells among dorsal *olig2*^+^ cells) at 5 dpf. (**G**) Graphical representation of the progression of maturing dorsal *mbp*^+^ OLs thoughout development. Scale bar = 20 μm. Data indicate means ± S.E.M, and dots represent individual larvae. *p<0.05; Statistical significance was drawn by ordinary one-way ANOVA followed by Sidak’s *post-hoc* and mixed-effects analysis followed by Tukey’s *post-hoc.* n^72^ ^hpf^ = 6-10 larvae per condition; n^5^ ^dpf^ = 5-12 larvae per condition.

At this developmental stage, Aβo-injected larvae exhibited an excess of *mbp*^+^ cells in the dorsal spinal cord compared to vehicle-injected larvae (**Figure 4E**), with no changes in *olig2*^+^ (**Figure 4B, 4D**) or total *mbp*^+^ cells (**Figure 4C**). Once again, Gö6983 treatment inhibited the effect triggered by Aβo (vehicle 9.58 ± 1.13 cells vs Aβo 14.25 ± 1.5 cells vs Aβo + Gö6983 7.4 ± 1.4 cells) (**Figure 4E**).

Regarding the proportion of myelinating OLs, consistent with our hypothesis, only 36% of dorsal OLs were *mbp*^+^ in the control larvae, compared to 53% in Aβo-treated larvae and 33% when PKC was inhibited (**Figure 4F**). Although no remarkable changes were observed in the number of mature myelinating OLs at 72 hpf (**Supplementary figure 5**), this transgenic line enabled us to conclude that OL maturation is significantly promoted by Aβo at 5 dpf, effect that was effectively reversed by PKC inhibitor Gö6983 (**Figure 4G**).

Taken together, these data suggest that Aβo induce precocious differentiation, resulting in altered OL maturation. Furthermore, Gö698 effectively inhibits these effects, indicating the involvement of the PKC signaling pathway in the Aβo-induced misregulation of OL differentiation and maturation.

### Aβo induce myelin excess in the dorsal spinal cord through PKC

Next, we investigated whether Aβo-induced OL differentiation and maturation could affect myelination *in vivo*. To assess myelin sheaths in individual OLs, we transiently expressed *mbpa*:EGFP-CAAX by microinjection into 1-cell stage zebrafish embryos. This genetic construct leads to the expression of membrane-tethered EGFP in the myelin tracts of the larvae. Aβo or vehicle injections were administered followed by Gö6983 treatments, and myelin sheath number per cell and sheath length were measured in the dorsal spinal cord at 5 dpf.

Aβo significantly increased the number of myelin sheaths per cell by around 40% in the dorsal spinal cord compared to vehicle-injected animals. Interestingly, while PKC inhibition alone did not significantly alter sheath numbers, treatment of Aβo-injected zebrafish larvae with Gö6983 restored the aforementioned Aβo-induced increase to control levels (vehicle 9.08 ± 0.9 sheaths; Aβo 12.7 ± 0.9 sheaths; Gö6987 11.1 ± 0.73 sheaths; Aβo + Gö6983 9.38 ± 0.79 sheaths) (**Figure 5A, B**). This change in sheath number per cell occurred without any alteration in myelin sheath length (**Figure 5C**).

**Figure 5.**
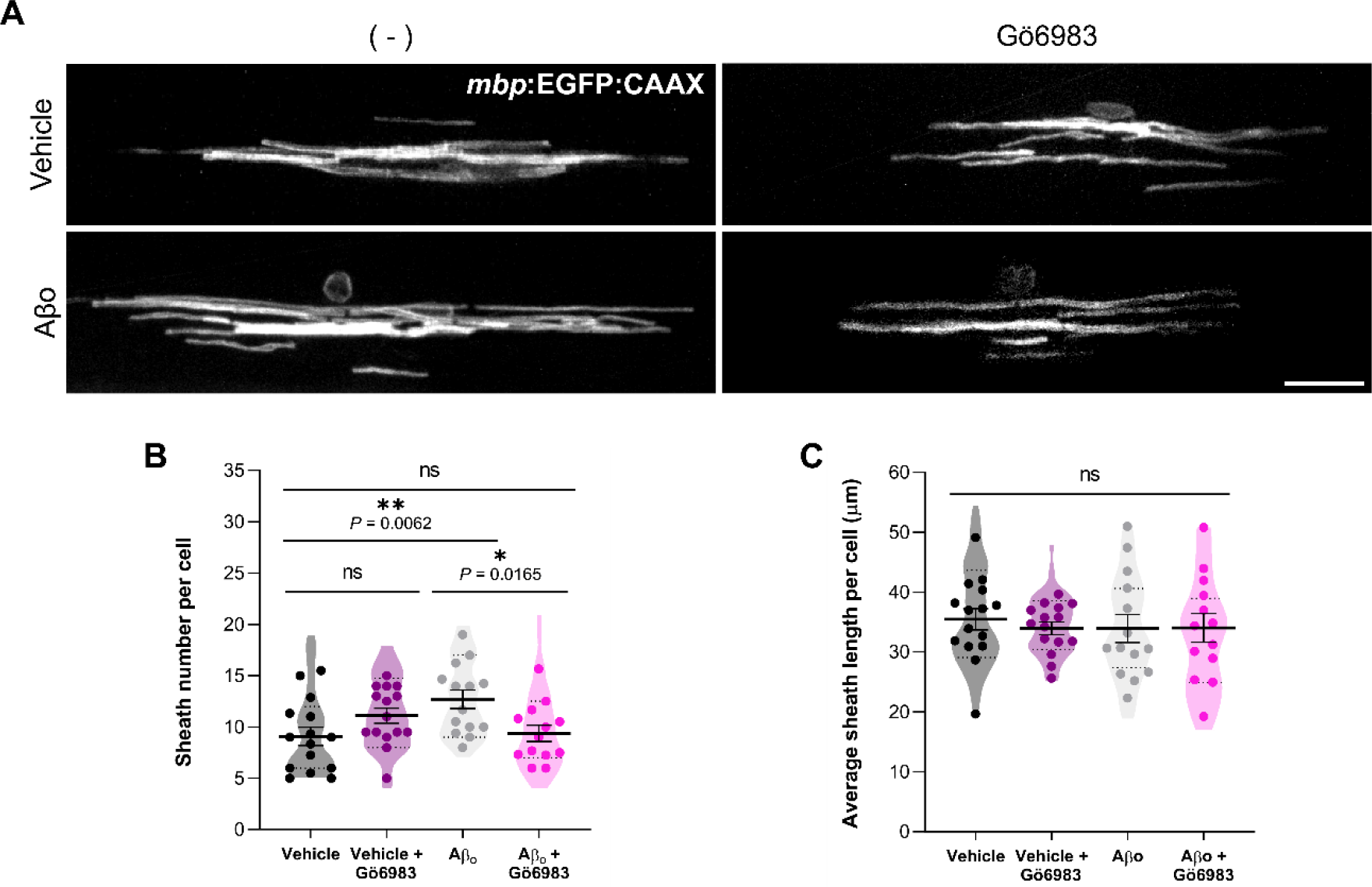
Aβo dysregulate dorsal myelin sheath number per oligodendrocyte via PKC. The *mbpa*:EGFP-CAAX plasmid was injected into zebrafish embryos at the 1-cell stage, followed by the administration of 10 µM of Aβo or its vehicle into the hindbrain ventricle of zebrafish larvae at 24 hpf. 500 nM Gö6983 treatments were performed at 48 hpf, and live imaging of myelin sheaths was performed at 5 dpf. (**A**) Representative fluorescent images of OLs in the spinal cord of live zebrafish larvae. (**B**) Histograms illustrating the sheath number per cell and (**C**) the average individual sheath length. Scale bar = 20 μm. Data are presented as means ± S.E.M, and dots represent individual larvae. *p<0.05, p<**0.01; Statistical significance was determined by two-way ANOVA followed by Sidak’s *post-hoc*.

Subsequently, we wondered whether the reported increase in the numbers of dorsal myelin sheaths per cell could result in notable changes in myelination. To address this question, electron microscopy was conducted on 8 dpf zebrafish larvae, and the myelinated axons within the dorsal region of the spinal cord were analyzed (**Figure 6A, B**).

**Figure 6.**
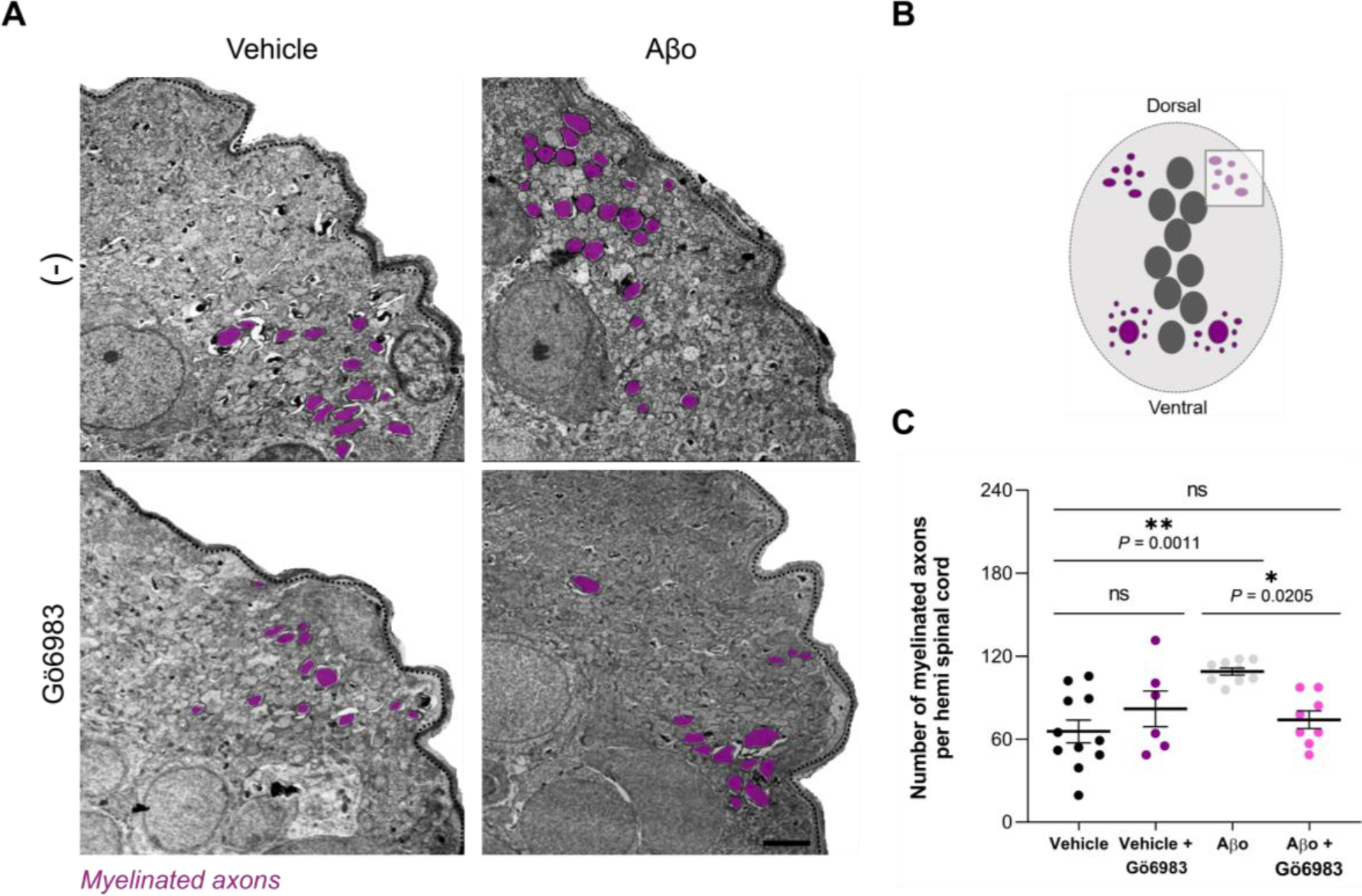
Aβo increase the number of dorsal myelinated axons through PKC. Zebrafish larvae were intracerebroventricularly injected with Aβo (10 µM) or its vehicle at 24 hpf, and some were exposed to Gö6983 (500 nM) at 48 hpf. Larvae were fixed for electron microscopy analysis at 8 dpf. (**A**) Representative electron micrographs of the dorsal spinal cord of 8 dpf zebrafish larvae, with myelinated axons shaded in purple. (**B**) Schematic illustration of a cross-section of the zebrafish spinal cord. The square represents the analyzed area. (**C**) Histogram showing the number of myelinated axons in the dorsal area for each condition. Scale bar = 2 µm. Data indicate means ± S.E.M, with dots representing individual larvae. *p<0.05, p<**0.01; Statistical significance was determined by two-way ANOVA followed by Sidak’s *post-hoc*.

In line with our previous findings, a significant increase of approximately 66% in the number of dorsal myelinated axons was observed in Aβo-injected zebrafish larvae compared to vehicle-injected larvae (108.9 ± 2.64 and 65.65 ± 8.24 myelinated axons, respectively). Moreover, while Gö6983 alone had minimal non-significant impact on dorsal myelination (81.98 ± 12.96 myelinated axons), PKC inhibition in Aβ-injected larvae nearly completely reversed the myelin excess induced by Aβo (74.07 ± 6.44 myelinated axons) (**Figure 6C**).

Altogether, we demonstrated that Aβo induce precocious OL differentiation leading to increased myelination, and we identified PKC as a key player in Aβo-induced dysregulation *in vivo*.

## Discussion

Growing evidence suggests that OL and myelin damage play significant roles in AD, with myelin impairment potentially leading to neuronal dysfunction and cognitive decline. However, the potential effects of Aβo on OL differentiation and myelination *in vivo* remain poorly understood. In recent years, zebrafish have emerged as a well-established model organism for studying OL and myelin biology *in vivo*, thanks to their optical transparency during embryonic and larval stages, which enables real-time observation of OL development, myelin formation, and myelin dynamics. Additionally, the rapid external development of the embryos and their genetic tractability facilitate genetic manipulation and screening approaches, allowing for the study of specific genes and signaling pathways involved in myelination. Importantly, many aspects of OL biology and myelin formation are highly conserved between zebrafish and mammals, making zebrafish an attractive model for investigating fundamental principles relevant to human health and disease (Choi et al., 2021; Preston & Macklin, 2015).

In this study, we investigated the effects of ICV injection of Aβo on OLs and myelin in the developing spinal cord of zebrafish larvae. For that, we focused on the expression of two well-established genes known to play pivotal roles in OL differentiation and myelination: *myrf*, primarily expressed in differentiating OLs, and *mbp*, expressed in mature myelinating OLs. Our analysis revealed that Aβo administration leads to alterations in the expression levels of both *myrf* and *mbp* mRNAs during larval development. Initially, *myrf* expression was significantly elevated in Aβo-injected larvae but decreased by 5 dpf. Similarly, consistent with *myrf, mbp* expression showed a notable increase in the presence of Aβo at early developmental stages, which was then normalized to control levels as development progressed. This finding was surprising, considering that myelination typically commences around 72 hpf (Buckley et al., 2010) and *mbp* expression is usually barely detectable at 48 hpf, as confirmed by the vehicle-injected zebrafish. The early induction of *mbp* mRNA observed in Aβo-injected larvae at 48 hpf suggested an accelerated myelination process in the presence of Aβo. Furthermore, we also identified *mbp* transcript accumulation in the dorsal region of the spinal cord, indicating enhanced OL differentiation and migration. Consistently, using live imaging of transgenic reporter lines for *olig2, myrf* and *mbp* transcripts, we found that Aβo exposure increased the number of *myrf*^+^ and *mbp*^+^ OLs in the dorsal spinal cord at 72 hpf and 5 dpf, respectively, without altering the total cell count. In line with these results, several studies have reported increased oligodendrogenesis in early stages of AD mouse models (Desai et al., 2010; Ferreira et al., 2020).These results collectively suggest a precocious differentiation and maturation of OLs in the presence of Aβo, without affecting cell proliferation or viability.

Accordingly, we also demonstrated that Aβo increase the number of myelin sheaths per OL in the dorsal spinal cord of zebrafish larvae, with no changes in sheath length, indicating that the observed transitory changes in transcription factor mRNA levels were sufficient to induce significant changes in myelination. Interestingly, this result is consistent with findings from a previous study where a transgenic zebrafish line with constitutively active Fyn kinase in all myelinating OLs showed an increase in sheath number per cell, without alterations in OL number or myelin sheath length (Czopka et al., 2013), which is comparable to what we observed in Aβo-injected zebrafish larvae. This suggests a potential involvement of Fyn kinase in mediating the effects of Aβo on myelination *in vivo*. Interestingly, Fyn has previously been described as a downstream target of Aβ both *in vitro* and in animal models (Boehm, 2013; Quintela-López et al., 2019). Furthermore, we observed elevated numbers of myelinated axons in the dorsal spinal cord of the zebrafish larvae exposed to Aβo. Similarly, an increase in myelin formation has also been described in the APP/PS1 AD mouse model (Chen et al., 2021). However, increased myelination does not necessarily indicate a beneficial outcome. The precise and coordinated production of myelin is essential for the correct development and functioning of the CNS. Indeed, multiple studies have reported myelin abnormalities in AD mouse models, including 3xTg-AD and APP/PS1 mice (Behrendt et al., 2013; Desai et al., 2009, 2010). More interestingly, previous zebrafish studies have shown that Aβo-injected zebrafish exhibit reduced avoidance of aversive stimuli compared to animals injected with vehicle alone (Nery et al., 2014). Consequently, it is plausible to speculate that the hypermyelination resulting from precociously differentiated OLs in our Aβo-injected zebrafish larvae may be aberrant or dysfunctional; however, further investigations are needed to confirm this hypothesis. These results together highlight the relevance of maintaining appropriate transcription factor levels and OL differentiation timing during development.

Aβo are promiscuous molecules able to signal through a repertoire of receptors and signaling pathways, promoting a wide range of effects in neurons, but also in OLs (Viola & Klein, 2015). Therefore, elucidating the underlying molecular mechanisms of Aβo is both of the utmost importance and challenging. In this study, we focused on PKC as a candidate kinase to modulate the Aβo-induced signaling pathway leading OL and myelin alterations. The diversity of PKC isoforms and the range of available inhibitors add complexity to this field of study, resulting in discrepancies in the literature. Some researchers report therapeutic effects of PKC activation in AD models (Etcheberrigaray et al., 2004), while others show that specific inhibition of PKCδ reverses AD phenotypes (Du et al., 2018). Nevertheless, the impact of PKC on OL and myelin dysfunctions has not been addressed. Here, we demonstrated that PKC inhibition with the pan-PKC inhibitor Gö6983 effectively reverses both Aβo-induced precocious OL differentiation and maturation, as well as the resulting hypermyelination. Importantly, this outcome not only confirms PKC’s involvement in these biological processes and the Aβo-induced signaling, but it also highlights Gö6983’s efficacy in mitigating OL and myelin alterations induced by Aβo in the developing spinal cord of zebrafish larvae. Interestingly, regarding the previously mentioned role of Fyn kinase in myelin modulation (Czopka et al., 2013), our findings suggest that PKC inhibition counteracts the effects of Fyn kinase on myelin, indicating a potential common pathway for both kinases. However, whether PKC acts upstream or downstream of Fyn remains to be elucidated. In summary, our findings strongly indicate that Aβo disrupt OL differentiation and myelination processes *in vivo* via a signaling pathway involving PKC activity.

## Acknowledgements

We thank C. Kearns, O. A. Mendez, and C. A. Doll for technical assistance and advice. We also thank the SGIKER platform from the University of the Basque Country (UPV/EHU) for the technical assistance with the electron microscopy.

## Author’s contributions

UB and A G-B designed and performed the experiments, analyzed and interpreted data, and wrote the manuscript. L B-C and MV S performed the experiments, analyzed and interpreted data. JL Z provided help to design the project, with data interpretation and reviewed the manuscript. EA and BA contributed to the conception and design of the project, analyzed and interpreted data, and wrote the paper. All authors have read and approved the final manuscript.

## Supplementary Material

### Supplementary methods

#### Intracerebroventricular injection of dextran in zebrafish larvae

At 24 hpf, all zebrafish embryos had their chorions removed for the brain ventricle injection procedure following previously established protocols (Gutzman & Sive, 2009; Nery et al., 2014). Embryos were anesthetized with Tricaine (Sigma-Aldrich) and microinjected in the hindbrain ventricle with 5–10 nl of a fresh injection buffer containing 10% fluorescein-labeled dextran (Invitrogen) and 10% Phenol Red in 0.4 M KCl in nuclease-free water. Subsequently, injected zebrafish were fixed in 4% PFA in PBS, gently rocking O/N at 4 °C, and dextran spreading was visualized both in the brain and the spinal cord using a Zeiss Cell Observer 2D 25 Spinning Disk confocal system (Carl Zeiss Microscopy) with a 40X oil-immersion objective.

#### Cortical oligodendrocyte culture

Highly enriched OPCs were prepared from mixed glial cultures obtained from newborn (P0–P2) Sprague–Dawley rat forebrain cortices as previously described (McCarthy & De Vellis, 1980) with minor modifications (Sánchez-Gómez et al., 2018). Briefly, forebrains were removed from the skulls and the cortices were enzymatically and mechanically digested. Then, tissue was plated in Iscove’s Modified Dulbecco’s Medium supplemented with 10% fetal bovine serum. The mixed glial cells were grown in poly-D-lysine (PDL) treated T75 flasks until they were confluent (8–10 days). Microglia were separated from the cultures by shaking the flasks on a rotary shaker. OPCs were isolated by additional shaking for 18 h (200 rpm, 37°C, 5% CO2). OPCs were seeded on to PDL-coated coverslips at densities of 8x10^4^ cells/well and were maintained at 37 °C and 5% CO2 for 3 days in a chemically defined differentiation SATO medium.

#### Western blot

OLs were exposed to Aβ peptides as indicated. Cells were scraped in sodium dodecyl sulfate (SDS) sample buffer on ice to enhance the lysis process and avoid protein degradation. Samples were boiled at 95 °C for 8 min, size-separated in 4-20% polyacrylamide-SDS Criterion TGX Precast gels (Bio-Rad) and transferred to Trans-Blot Turbo Midi Nitrocellulose Transfer Packs (Bio-Rad). Membranes were blocked in 5% BSA in Tris-buffered saline/0.05% Tween-20 and proteins were detected by specific primary antibodies against PKC (#ab179521, 1:1000; Abcam) and p-PKC (#9371, 1:1000; Cell Signaling). Membranes were incubated with horseradish peroxidase-conjugated secondary antibodies (1:5000). The protein bands were detected with a ChemiDoc XRS Imaging System (Bio-Rad), and quantified by using Image Lab 6.0.1 (Bio-Rad) software.

## Supplementary figures

**Supplementary figure 1.**
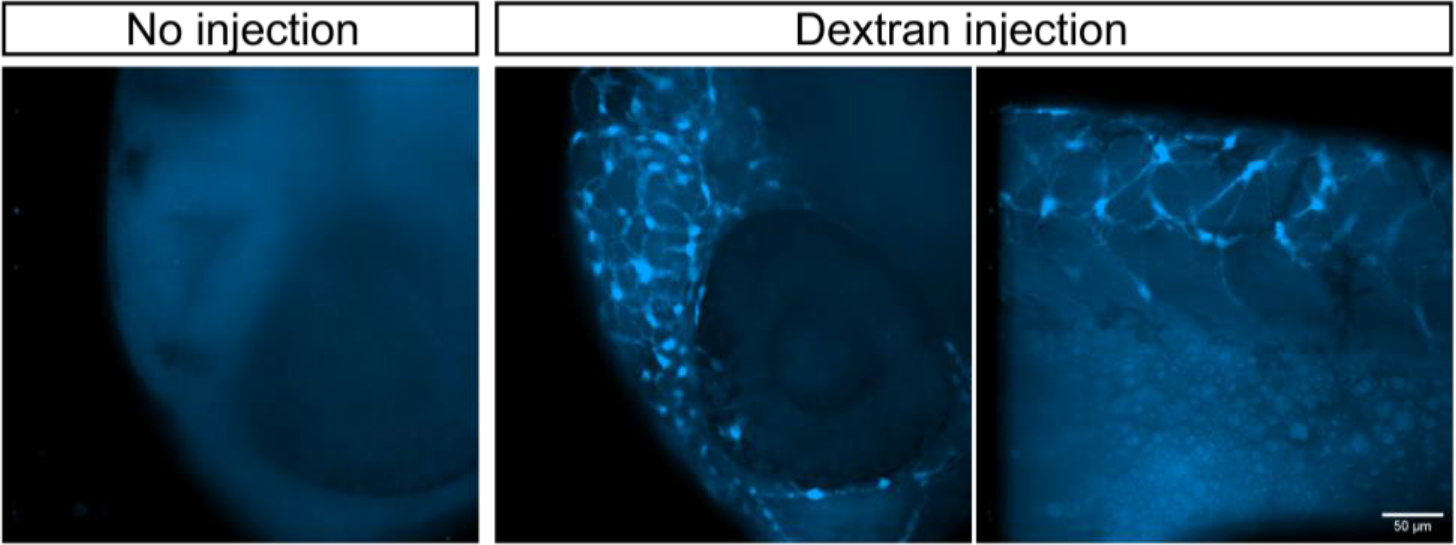
Intraventricular injection of fluorescently-labeled dextran into 24 hpf zebrafish larvae.

**Supplementary figure 2.**
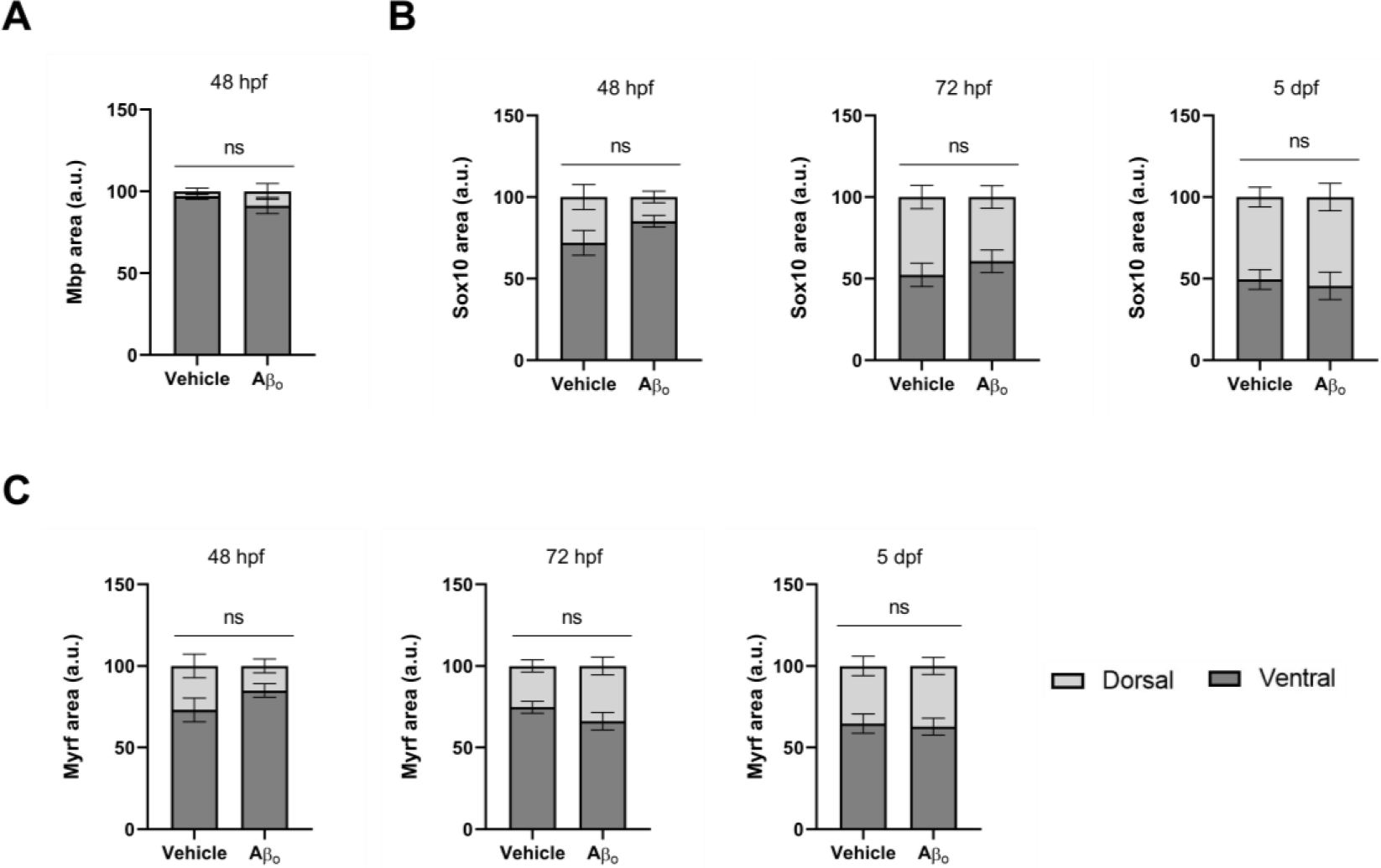
Regional distribution of (A) *mbp*, (B) *sox10* and (C) *myrf* mRNA in the developing zebrafish spinal cord. Histograms show percentages of dorsal and ventral distribution relative to total area. Statistical significance was determined by two-way ANOVA followed by Sidak’s post-hoc test. n = 9-11 larvae per condition and time-point.

**Supplementary figure 3.**
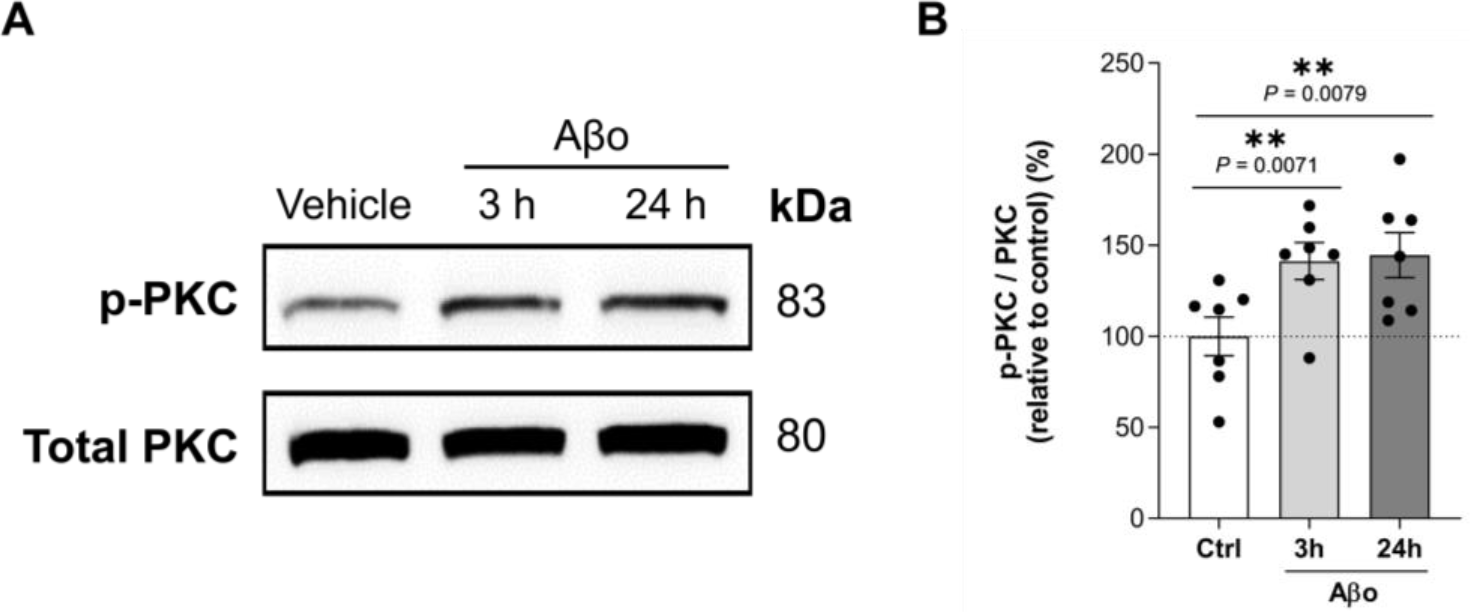
Aβo activate PKC in primary cultured oligodendrocytes. (**A**) Western blot analysis and (**B**) quantification of PKC phosphorylation levels in control and Aβo-treated OLs. Histogram represent protein expression levels as percentages (%) relative to control cells. Data are presented as means ± S.E.M; dots represent individual experiments and violin plot represents quantification of individual cells. **p<0.01; Statistical significance was determined by one-way ANOVA followed by Dunnett’s *post-hoc* test.

**Supplementary figure 4.**
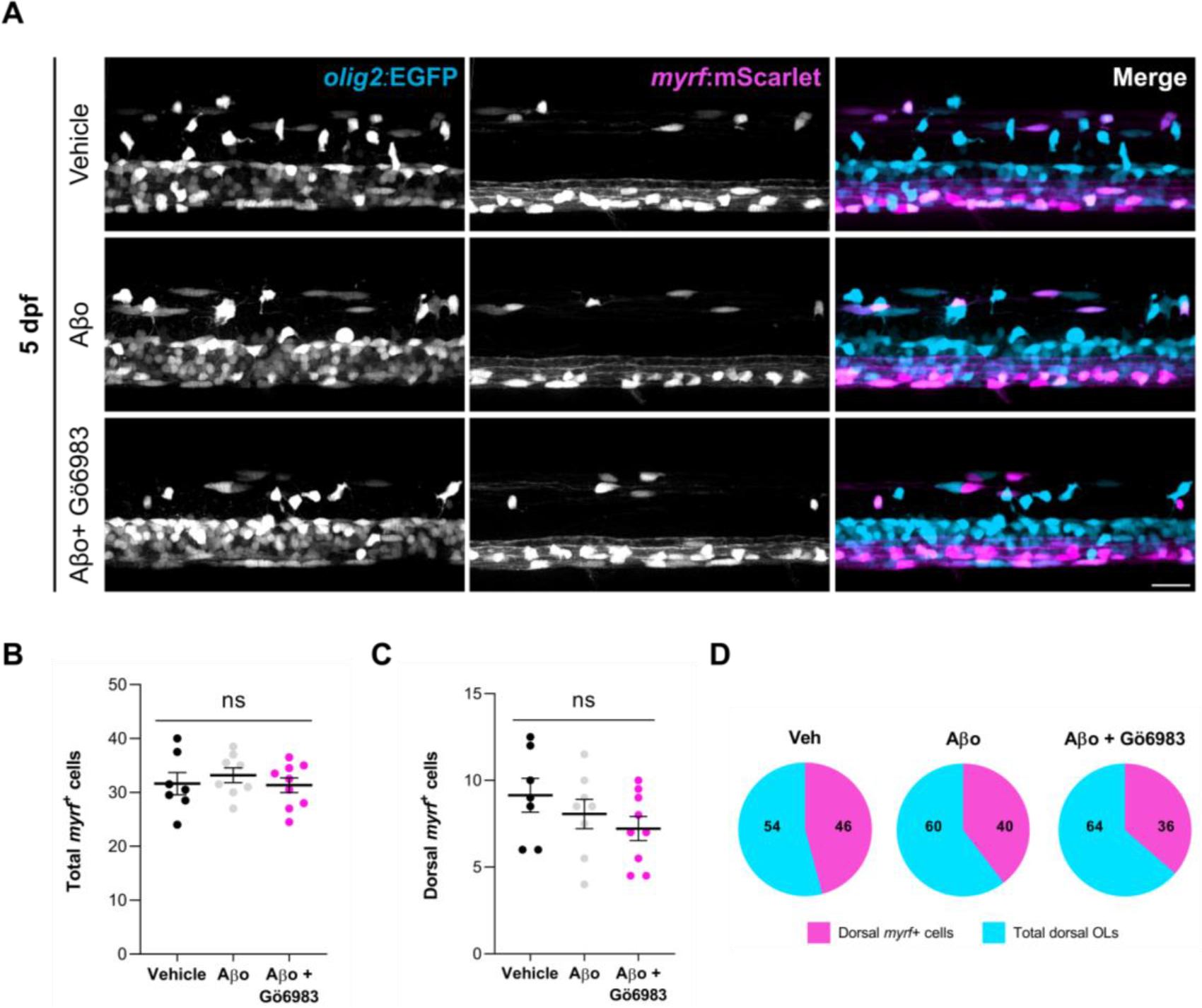
***Myrf*^+^ cell numbers are unchanged at 5 dpf.** (**A**) Representative lateral images of the spinal cord of live transgenic larvae stably expressing *olig2*:EGFP and *myrf*:mScarlet, at 5 dpf. Graphs showing the number of (**B**) total and (**C**) dorsal *myrf*^+^ cells. (**D**) Pie charts showing the ratio of differentiating dorsal OLs (percentage of dorsal *myrf*^+^ cells among dorsal *olig2*^+^ cells) at 5 dpf for each condition. Scale bar = 20 μm. Data indicate means ± S.E.M, and dots represent individual larvae. Statistical significance was determined by ordinary one-way ANOVA followed by Sidak’s *post-hoc*.

**Supplementary figure 5.**
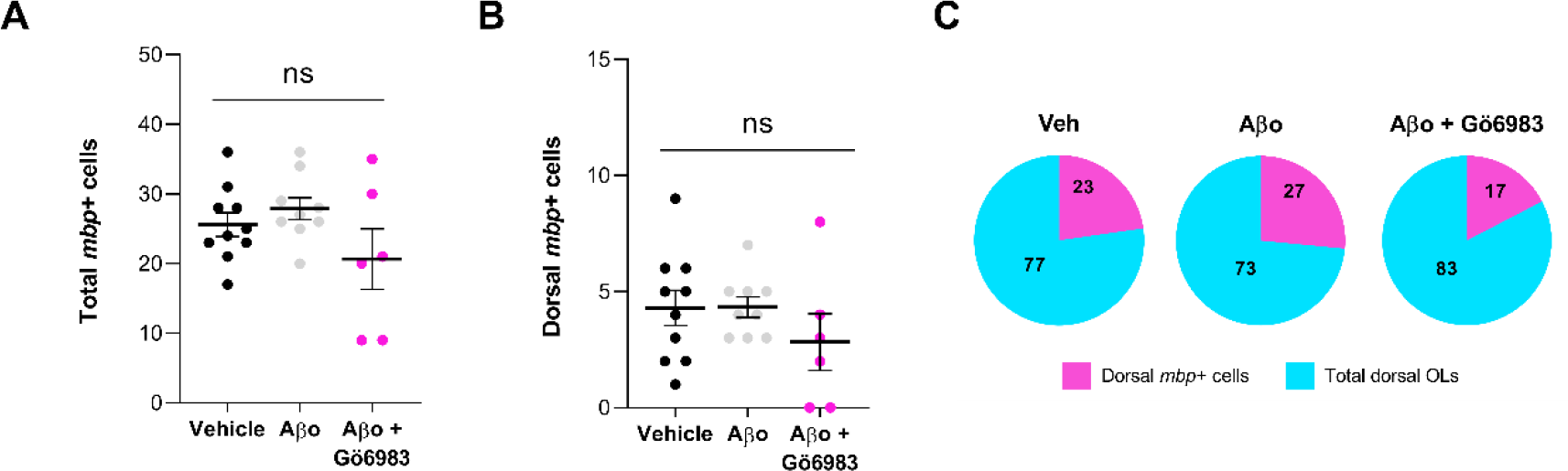
***Mbp*^+^ cell numbers are unchanged at 72 hpf.** Graphs showing the number of (**A**) total and (**B**) dorsal *mbp*^+^ cells in the spinal cord of live transgenic larvae stably expressing *olig2*:EGFP and *mbpa*:tagRFPT, at 72 hpf. (**C**) Pie charts showing the ratio of differentiating dorsal OLs (percentage of dorsal *mbp*^+^ cells among dorsal *olig2*^+^ cells) at 72 hpf for each condition. Data indicate means ± S.E.M, and dots represent individual larvae. Statistical significance was determined by ordinary one-way ANOVA followed by Sidak’s *post-hoc*.

